# Proteomic profiling of mitochondrial-derived vesicles in brain reveals enrichment of respiratory complex sub-assemblies and small TIM chaperones

**DOI:** 10.1101/2020.07.06.189993

**Authors:** Rosalind F. Roberts, Andrew N. Bayne, Thomas Goiran, Dominique Lévesque, François-Michel Boisvert, Jean-François Trempe, Edward A. Fon

## Abstract

The generation of mitochondrial-derived vesicles (MDVs) is implicated in a plethora of vital cell functions, from mitochondrial quality control to peroxisomal biogenesis. The discovery of distinct subtypes of MDVs has revealed the selective inclusion of mitochondrial cargo in response to varying stimuli. However, the true scope and variety of MDVs is currently unclear, and unbiased approaches have yet to be used to understand their biology. Furthermore, as mitochondrial dysfunction has been implicated in many neurodegenerative diseases, it is essential to understand MDV pathways in the nervous system. To address this, we sought to identify the cargo in brain MDVs. We used an in vitro budding assay and proteomic approach to identify proteins selectively enriched in MDVs. 72 proteins were identified as MDV enriched, of which 31% were OXPHOS proteins. Interestingly, the OXPHOS proteins localized to specific modules of the respiratory complexes, hinting at the inclusion of sub-assemblies in MDVs. Small TIM chaperones were also highly enriched in MDVs, linking mitochondrial chaperone-mediated protein transport to MDV formation. As the two Parkinson’s disease genes PINK1 and Parkin have been previously implicated in MDV biogenesis in response to oxidative stress, we compared the MDV proteomes from the brains of wild-type mice with those of PINK1^-/-^ and Parkin^-/-^ mice. No significant difference was found, suggesting that PINK1- and Parkin-dependent MDVs make up a small proportion of all MDVs in the brain. Our findings demonstrate a previously uncovered landscape of MDV complexity and provide a foundation from which to discover further novel MDV functions.

## Introduction

Mitochondrial-derived vesicles (MDVs) are a recently described biological phenomenon, which were initially reported as Drp1-independent vesicles with a 70-100 nm diameter^1^. They mediate communication between mitochondria and other organelles, as well as external immune cells, via vesicular trafficking of mitochondrial material^2^ MDVs have been implicated in processes as diverse as mitochondrial quality control, peroxisomal biogenesis and antigen presentation, highlighting their crucial role in linking dynamic mitochondrial function to the wider cellular ecosystem^3^. As a result, dysfunction of MDV formation or trafficking may be a potential mechanism in diseases of many tissues, particularly those with high dependence on mitochondrial function, such as the brain.

Cargo selectivity is a fundamental feature of MDVs, suggesting intricate mechanisms for selection and packaging of specific cargo. Distinct MDV subtypes are believed to incorporate distinct cargo profiles, which are likely related to their function. For example, MDVs containing Pex3 and Pex14 are crucial for peroxisomal biogenesis and fuse with ER-derived vesicles containing Pex16 to form the mature organelle^4^ Pex3/Pex14 MDVs are distinct from MAPL-positive MDVs, which also traffic to the peroxisome but the function of which remain unknown^1, 5^. Furthermore, the Parkinson’s disease linked proteins PINK1 and Parkin regulate the formation of several classes of MDVs, suggesting there may be multiple layers of MDV biogenesis regulation with each responding to different stimuli or in a cell-type specific manner. For example, PINK1 and Parkin mediate the formation of PDH-positive/TOM20-negative MDVs in response to oxidative stress, which traffic to the lysosome for turnover^6^. TOM20-positive/PDH-negative MDVs, which also traffic to the lysosome, are not dependent on PINK1 and Parkin for their biogenesis^6–17^, although Parkin may be required for their endosomal delivery^8^. In macrophages, bacterial infection triggers the PINK1/Parkin-dependent formation of Sod2-positive/Tom20 negative MDVs that traffic to the phagosome to generate anti-microbial hydrogen peroxide^9^ Inhibition of lysosomal degradation increases their phagosomal targeting, suggesting this class of MDV may have dual targets^9^ Intriguingly, heat shock or lipopolysaccharide treatment of macrophages leads to the trafficking of MDVs containing mitochondrial antigens (OGDH) for presentation on MHC class I molecules, in a process that is negatively regulated by PINK1 and Parkin^10–11^. Therefore, a complex picture of MDV regulation is beginning to emerge.

While the number of MDV classes described expands, the full molecular details of the cargo and regulatory molecules for each MDV subtype remain relatively obscure. In particular, their study relies upon the arbitrary selection of positive and negative cargo to allow their visualization by immunofluorescence, often based upon the availability of suitable antibodies. The cargo profile of any one MDV class is usually restricted to one or two positive and negative markers, again limiting our understanding of the complexity of MDV biogenesis, regulation and trafficking.

To address this issue and reveal the complexity of MDV biology, an unbiased approach to identify highly enriched cargo is essential. Furthermore, given the vast diversity of mitochondria between tissues^12^, understanding the ‘MDVome’ with a tissue specific approach is key to unlocking their involvement in biology and disease. We therefore sought to obtain a full proteome of brain MDVs to gain further insight into their biology and to facilitate functional studies of MDVs in neurons and glia. Additionally, we attempted to identify MDV cargo dependent on PINK1 and Parkin, two Parkinson’s disease genes previously implicated in MDV biogenesis in response to oxidative stress, to further elucidate their role in mitochondrial quality control.

Here, we present the first unbiased proteomic analysis of total brain MDVs. Our study reveals the enrichment of OXPHOS and small TIM chaperone proteins in MDVs. As we were not able to identify specific components of PINK1/Parkin-dependent MDVs, we propose that this class of MDVs may form a small proportion of the total pool of MDVs. Therefore, the full biological diversity of MDVs may be much broader than previously anticipated. Our survey of the MDV proteome will allow the full spectrum of MDV function to be uncovered.

## Experimental procedures

### Animals

For all proteomic experiments, tissue from two or three mice was pooled for one biological replicate. Data set 1 comprises three biological replicates from PINK1^-/-^ mice and three biological replicates from their wild-type littermates. Data set 2 comprises three biological replicates from Parkin^-/-^ mice and three biological replicates from their wild-type littermates. All animal work was performed in accordance with the recommendations and guidelines of the Canadian Council on Animal Care (CCAC), the McGill University Animal Care Committee (UACC), and The Neuro Animal Care and Use Program.

### In vitro budding assay

Budding assays were performed as previously described with slight modifications (Soubannier et al, 2012; McLelland et al, 2017). Briefly 10-12 week old mice were anaesthetized with isofluorane and euthanized by exposure to carbon dioxide. The brains were swiftly extracted and placed in cold mitochondrial isolation buffer (MIB - 20 mM HEPES, pH 7.4, 220 mM mannitol, 68 mM sucrose, 76 mM KCl, 4 mM KOAc, 2 mM MgCl2 plus protease inhibitors added just before use). Brains were washed twice in cold MIB, then weighed and homogenized on ice in 6 ml MIB per g tissue using motor-driven PTFE tissue grinders at 1600 rpm (4 strokes with a loose pestle followed by 8 strokes with a tight pestle).

An aliquot of the resulting homogenate (H) was removed then the homogenate was centrifuged at 1,300 x *g* for 10 minutes at 4 °C. The supernatant was carefully removed and centrifuged at 21,000 x *g* for 10 minutes at 4 °C. The supernatant (S+LM) was removed and transferred to a new centrifuge tube and the pellet (HM) was resuspended in the same volume of MIB as the supernatant, and all were centrifuged at 21,000 x *g* for 10 minutes at 4 °C. The HM pellet was kept, resuspended in the same volume of MIB again and stored on ice. The supernatant from S+LM was carefully removed and centrifuged at 200,000 x *g* for 90 min at 4 °C. The supernatant was retained (S) and the pellet (LM) was washed twice with MIB, before resuspension in MIB with 2% Triton X-100.

Following protein quantification using BCA (Pierce), 4.5 mg HM was incubated with 1.125 mg S plus ATP regenerating mixture (1 mM ATP, 5 mM succinate, 80 μM ADP, 2 mM K2HPO4, pH 7.4) and 50 μM antimycin A or DMSO (0.1%). The volume was made up to 1 ml with MIB and the reaction was incubated at 37 °C for 1 hour, with vortexing after 30 minutes.

The reaction was stopped by centrifugation at 7,500 x *g* for 10 minutes at 4 °C. The supernatant containing MDVs was removed and mixed with 80% sucrose, to give a final sucrose percentage of 60%. This mixture was loaded at the bottom at a SW41 ultraclear tube (Beckman) and 50%, 40% and 30% sucrose were sequentially layered on top for a discontinuous sucrose gradient, which was centrifuged for 16 hours at 35,000 rpm (151,000 x *g* at *r_av_*) in SW41 rotor at 4 °C.

Fractions were removed as 0.5 ml or 1 ml aliquots and 25% of fractions from the 40-50% interface were retained for western blot analysis (5% loaded per lane). The remainder was stored at −80 °C until mass spectrometry preparation.

### Western blotting

Western blotting was performed using standard procedures. Briefly, 10% polyacrylamide gels were cast using the Bio-Rad mini system and samples were loaded following resuspension in 5x Laemmli buffer. For all blots shown, 3 μg of H, S, LM or HM were loaded and 5% of sucrose gradient fractions were loaded. Voltage of 100 V was applied for 30 minutes, followed by 150 V for 1 hour. Proteins were transferred to nitrocellulose membranes using the BioRad TurboBlot system, for 20 minutes at 25 V. All of the following steps were performed with gentle shaking. Membranes were blocked in 3% non-fat skim milk in PBS with 0.1% Tween for 1 hour at room temperature, followed by incubation with primary antibody in 1% milk in PBS with 0.1% Tween overnight at 4 °C. Following three washes in PBS with 0.1% Tween, membranes were incubated with the relevant secondary antibody diluted at 1:5000 in 1% milk in PBS with 0.1% Tween. Following three washes in PBS with 0.1% Tween, membranes were developed using Clarity Max (BioRad) in a ChemiDoc (BioRad).

### Antibodies

**Table 1:**
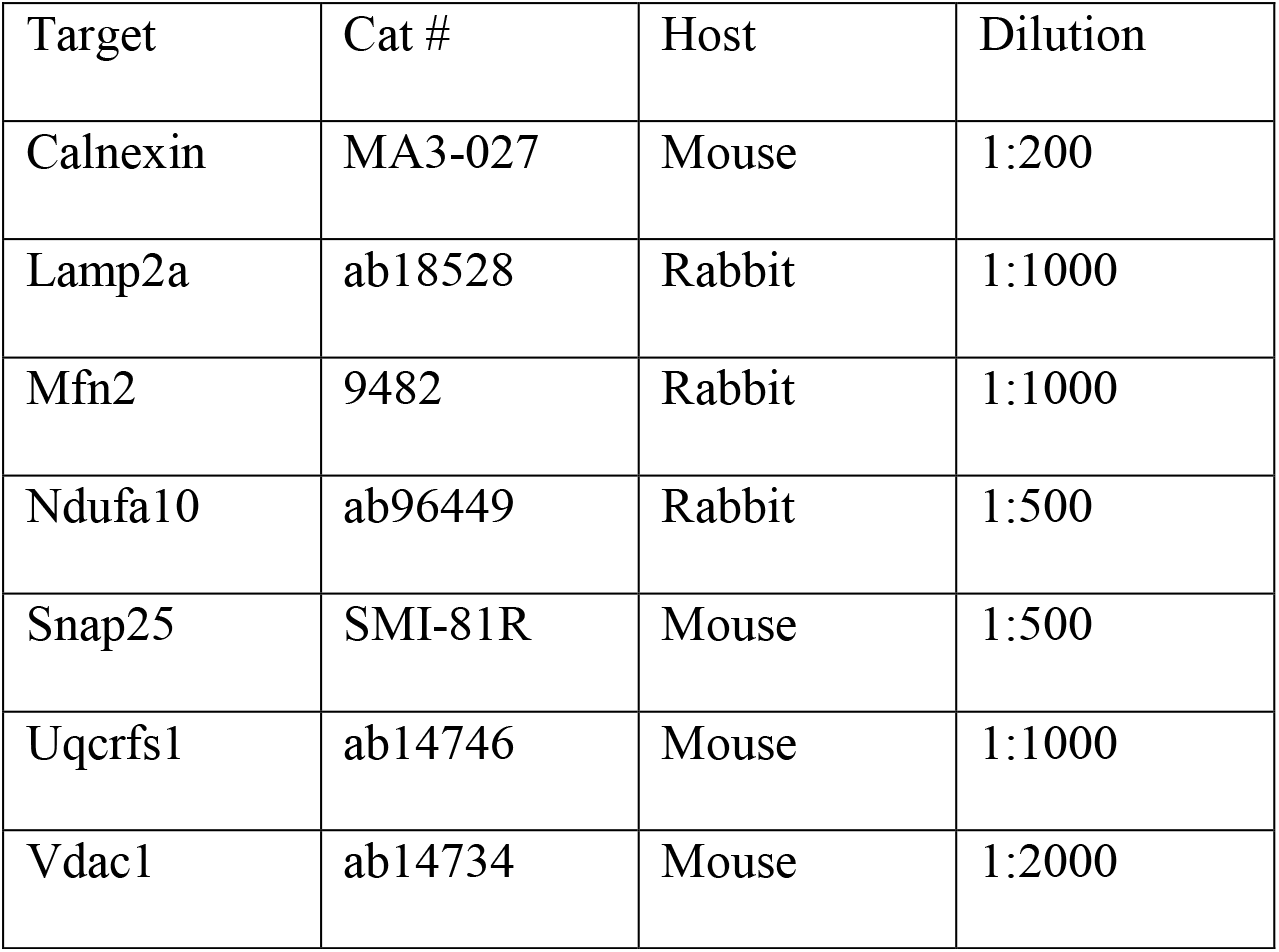
Antibodies used for western blotting in this study.

### Respiratory assay

Oxygen consumption rates were assessed using the MitoXpress respiratory assay (Agilent Technologies) and were performed according the manufacturer’s instructions. In a 96 well plate, 100 μg of HM/crude mitochondrial preparation was mixed with 10 μl 10X ATP regenerating mixture (10 mM ATP, 50 mM succinate, 800 μM ADP, 20 mM K_2_HPO_4_, pH 7.4) and volume made up to 90 μl with MIB. Immediately prior to loading the plate in the Perkin Elmer EnSpire plate reader pre-heated to 37 °C, 10 μl of MitoXpress reagent, and 1 μl of MIB, DMSO or antimycin A (final concentration: 50 μM) were added, followed by a layer of mineral oil. The plate was read with dual-TRF every 2 minutes for 90 minutes. The fluorescence intensity was plotted against time and the slope over the first 14 minutes was used to calculate the oxygen consumption rate (OCR) basally or following treatment with DMSO or antimycin A.

### Transmission electron microscopy

Sucrose gradient fractions at the 40/50% interface were diluted 1:1 with MIB and centrifuged at 200,000 x g for 1 hour at 4 degrees. The pellet was washed gently using a manual pipette twice with MIB, then dried for 10 minutes before resuspension in 20 μl of 20 mM HEPES. 5 μl of this preparation was applied to a copper formvar grid and left to dry for 10 minutes at room temperature. The grid was dipped in water followed by 2% uranyl acetate for 45 seconds. The excess was removed using filter paper and the grid left to dry for 2 hours at room temperature. The data were collected on a FM Spirit Tecnai electron microscope located at the Facility for Electron Microscopy Research at McGill University.

### Mass spectrometry sample preparation

For MDV sample preparation, sucrose gradient fractions at the 40/50% interface (boxed in red in Figure 1C) from the budding assay described above were diluted 1:1 with MIB and centrifuged at 200,000 x g for 1 hour at 4 degrees. The pellet was washed gently using a manual pipette twice with MIB, then dried for 10 minutes before resuspension in 20 μL of denaturing buffer (8 M urea, 1mM EDTA, 50mM TEAB pH8.5). Protein quantification using BCA (Pierce) showed that MDV samples were very low in abundance and the yield was variable. We determined in previous experiments that the average protein concentration at the 40/50% interface was approximately 500 ng. Therefore, we digested all MDV samples based on 500 ng protein, since we did not have enough material to perform a BCA following each budding assay.

**Figure 1.**
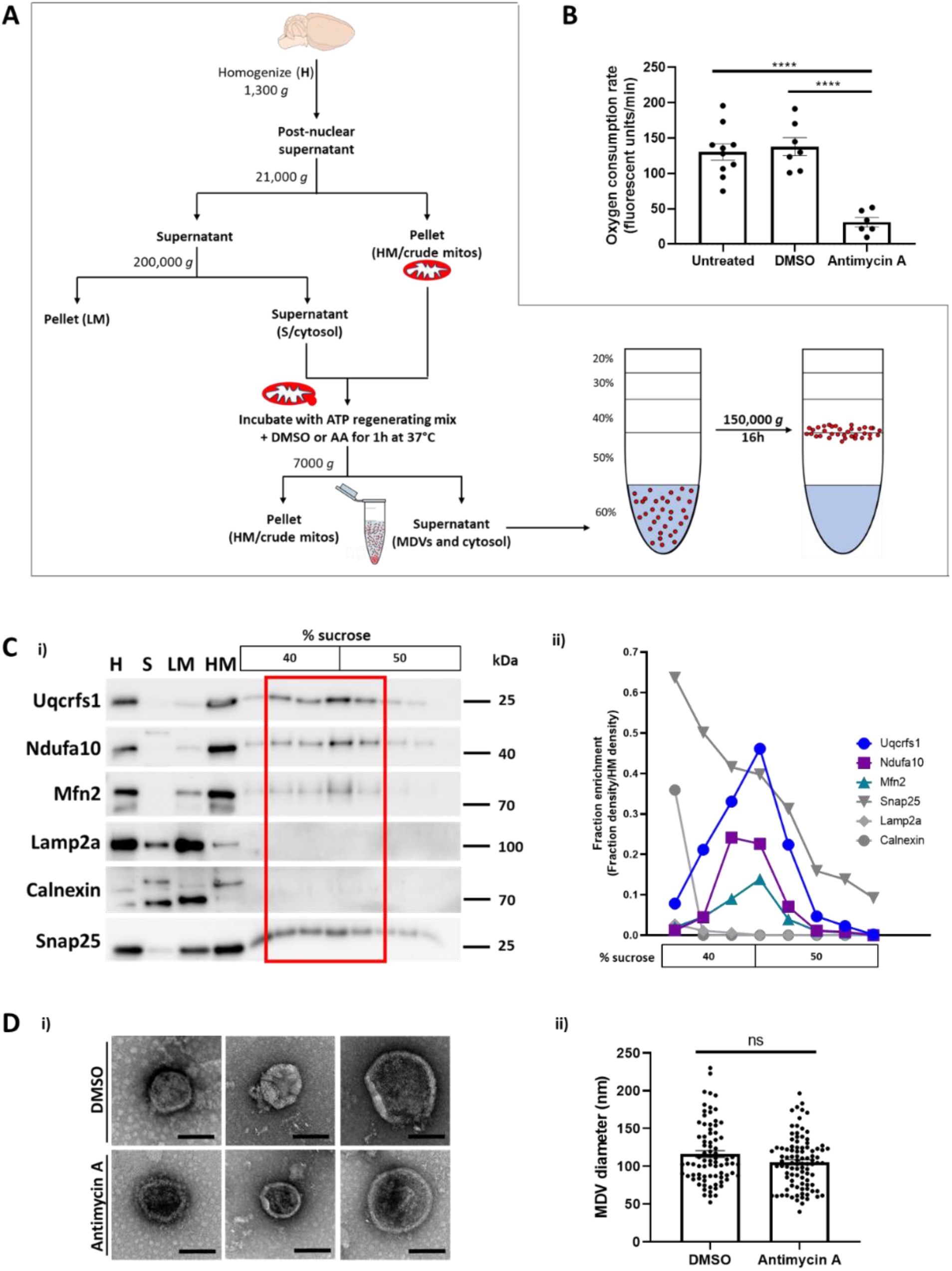
*In vitro* MDV budding assay in mouse brain. A) Schematic of *in vitro* MDV budding assay and purification of MDVs using sucrose gradients. B) Mitochondria (HM) isolated from wild-type mouse brain were metabolically active and responded to the complex III inhibitor antimycin A (AA), as assessed by MitoXpress respiratory assay. One way ANOVA with Tukey’s multiple comparisons test. ****: p<0.0001 C) MDVs were budded from mitochondria (HM) isolated from wild-type mouse brain in the presence of cytosol and 0.1% DMSO. Budding supernatants were separated on a discontinuous sucrose gradient and MDVs floated to the 40/50% interface (i). Mitochondrial proteins including Uqcrfs1, Ndufa10 and Mfn2 were enriched specifically at the 40/50% interface, while lysosomal (Lamp2a), ER (calnexin) and synaptic (Snap25) proteins were either not present or not enriched at this interface (ii). Blots and graph are representative of 2-5 independent experiments (Uqcrfs1, Ndufa10, Mfn2 – N=5; Snap 25 – N=2; Lamp2a and Calnexin – N=3). Error bars are not shown for clarity and can be found in the complementary Figure S2A. D) Transmission electron micrographs of purified MDVs stained with uranyl acetate. Scale bar: 100 nm. No significant difference in the diameter of MDVs between experimental conditions was found (unpaired two-tailed t-test).

For mitochondrial sample preparation, 50 μg of parental mitochondria (HM) were diluted in 1 mL MIB and centrifuged at 7,500 x *g* for 10 minutes at at 4 °C. The pellet was resuspended in 20 μL of denaturing buffer (8 M urea, 1 mM EDTA, 50 mM TEAB pH 8.5). The excess amount of HM compared to MDV, which are approximately 500 ng per sample, was used to allow us to assess the enrichment of MDV cargo compared with as complete a mitochondrial proteome as possible.

For mitochondrial samples submitted to the sucrose gradient as a control for methionine oxidation control levels, 950 μg HM was loaded at the bottom of the sucrose gradient in 60% final concentration sucrose, using the same method as for MDVs in the budding assay. 50%, 40% and 30% sucrose were sequentially layered on top for a discontinuous sucrose gradient, which was centrifuged for 16 hours at 35,000 rpm (151,000 x *g* at *r_av_*) in SW41 rotor at 4 °C. The bottom sucrose gradient fraction (60% sucrose), containing mitochondria, was diluted 1:1 and centrifuged at 200,000 x g for 1 hour at 4 degrees. The pellet was washed gently using a manual pipette twice with MIB, then dried for 10 minutes before resuspension in 20 μL of denaturing buffer (8 M urea, 1mM EDTA, 50mM TEAB pH8.5). All mitochondria samples processed in this way were digested based on 500 ng protein, since they were similarly low in abundance to the MDV samples.

Samples in denaturing buffer were reduced with 2.5 mM TCEP for 1 hour at 37 °C, alkylated with 10 mM fresh iodoacetamide for 30 min at room temperature, then pre-digested with Lys-C (Wako) in a 1:100 (μg sample/μg protease) ratio for 2 hours at 37°C in the dark. After cooling to 25°C, samples were diluted with 50 mM TEAB pH 8.5 to reduce the urea concentration to < 1.5 M, and trypsin (Sigma, sequencing grade) was added in a 1:50 ratio to incubate for 16 hours at 25°C. To stop the digestion, samples were acidified to 0.5% formic acid and 5% acetonitrile. Samples were loaded onto Pierce C18 Spin Columns (ThermoFisher), washed with 5% acetonitrile/0.5% formic acid, and eluted in 50% acetonitrile/0.1% formic acid. Elutions were dried under ambient temperature in a SpeedVac (ThermoFisher SPD-1010), then the peptides were resuspended in 25 μl of 1% FA buffer. Peptides were assayed using a NanoDrop spectrophotometer (Thermo Fisher Scientific, Waltham, USA) and read at an absorbance of 205 nm. The peptides were transferred to a glass vial (Thermo Fisher Scientific, Waltham, USA) and stored at −20°C until analysis by mass spectrometry.

### LC-MS/MS analysis

For LC-MS/MS, 250 ng of each sample was were injected into an HPLC (nanoElute, Bruker Daltonics) and loaded onto a trap column with a constant flow of 4 μl/min (Acclaim PepMap100 C18 column, 0.3 mm id x 5 mm, Dionex Corporation) then eluted onto an analytical C18 Column (1.9 μm beads size, 75 μm x 25 cm, PepSep). Peptides were eluted over a 2-hour gradient of acetonitrile (5-37%) in 0.1% FA at 500 nL/min while being injected into a TimsTOF Pro ion mobility mass spectrometer equipped with a Captive Spray nano electrospray source (Bruker Daltonics). Data was acquired using data-dependent auto-MS/MS with a 100-1700 m/z mass range, with PASEF enabled with a number of PASEF scans set at 10 (1.27 seconds duty cycle) and a dynamic exclusion of 0.4 minute, m/z dependent isolation window and collision energy of 42.0 eV. The target intensity was set to 20,000, with an intensity threshold of 2,500.

The mass spectrometry proteomics data have been deposited to the ProteomeXchange Consortium via the PRIDE partner repository^13^ with the dataset identifier PXD020197.

Reviewer account details:

Username

Password: 2bLSWrxw

### Protein and modification identification by MaxQuant analysis

The raw mass spectra were analyzed using MaxQuant (version 1.6.10.43) with MaxQuant tims-DDA specific parameters already embedded and the Uniprot mouse database (UP000000589_10090, 55,398 entries). The settings used for the MaxQuant analysis were: 2 miscleavages were allowed; minimum peptide length was set to 5; fixed modification was carbamidomethylation on cysteine; enzymes were Trypsin (K/R not before P); variable modifications included in the analysis were methionine oxidation, tyrosine nitration, tyrosine nitrosylation, tyrosine hydroxylation and protein N-terminal acetylation. Label-free quantification (LFQ) was chosen using default settings, including a minimum ratio count of 2. Peptide spectral match and protein false discovery rates and site decoy fraction were set to 0.05.

The resulting gene names list and spectral counts were subsequently analyzed following normalization by expressing each LFQ value as a percentage of the total LFQ intensity for all proteins detected within that sample (% total LFQ). Data were then imported into Perseus (v1.6.10.50) for analysis and transformed to the logarithmic scale (base 2). Proteins positive for at least one of the “Reverse”, “Potential contaminant” or “Only detected by site” categories, or with valid values in fewer than 70% of samples were eliminated. Volcano plots were generated to assess enrichment of proteins in MDVs compared to mitochondria, or the effect on MDV enrichment of experimental conditions (antimycin A treatment or genotype) using multiple t-tests with permutation-based FDR of 0.05 with 250 randomizations and s0 of 0.1.

For global assessment of post-translation modifications, the total number of peptide counts for a particular modification were divided by the total peptide count in a sample, then multiplied by 100 to give a percentage.

### Functional annotation and gene ontology analysis

Protein hits were assigned a functional annotation following integration of information from a variety of resources including gene and protein information databases (MitoCarta^14^, GeneCards^15^, COMPARTMENTS^16^, NCBI^17^, Uniprot^18^) and primary literature. Proteins that did not clearly fit into one of the following categories (OXPHOS, metabolism, channels, chaperones, redox signalling, mitochondrial, cytoskeletal/trafficking, ribosomal) were classed as non-mitochondrial. Proteins that have only been shown to be localized to synapses, the plasma membrane or non-nervous tissues were included in the non-mitochondrial category to avoid skewing by potential contaminants.

Principal component analysis (PCA) was performed in Perseus (v1.6.10.50), with the data processed as described in the MS analysis section. Proteins with valid values in fewer than 100% of samples were eliminated from the analysis.

STRING analysis (version 11.0) was performed on the list of proteins enriched in MDVs using https://string-db.org/^19^ The network edges represent confidence in the interaction (line thickness indicate the strength of the supporting data). A minimum required interaction score of 0.4 (medium) was selected. Disconnected nodes were hidden.

Gene ontology analysis of the proteins enriched in MDVs was performed using g:Profiler^20^ using the following settings. Statistical domain scope: only annotated genes. Significance threshold: g:SCS threshold. User threshold: 0.05. Gene Ontology biological process was used for the analysis.

## Results

### Generating mitochondrial-derived vesicles from brain mitochondria using an in vitro budding assay

To analyze MDVs using proteomics, we required a method to purify at least several hundred nanograms of brain MDVs. To achieve this, we adapted a previously developed *in vitro* budding assay^7–21^ to generate and purify MDVs derived from brain mitochondria.

Mouse brains were homogenized (H) and fractionated into supernatant (S/cytosol), light membrane (LM) and heavy membrane (HM/crude mitochondrial extract) fractions (Figure 1A). To confirm the suitability of the isolated brain mitochondria for the budding assay, we assessed the respiratory activity of HM (crude mitochondrial extract) using an oxygen-sensitive fluorophore, MitoXpress (Figure 1B). Brain-derived HM had oxygen consumption rates (OCR) within the expected range^22^ Treatment with 50 μM antimycin A, a complex III inhibitor, ablated respiratory activity as expected, while the vehicle control (0.1% DMSO) had no effect. We did not observe a difference in respiratory activity or response to antimycin A in wild-type animals compared with their PINK1 or Parkin null littermates (Figure S1A).

To stimulate MDV budding, we incubated HM with cytosol (S) and a succinate-based ATP regenerating mixture for 1 hour (Figure 1A). Following sucrose gradient purification of the post-reaction supernatant, we probed the fractions for the mitochondrial proteins Uqcrfs1 (CIII Rieske), which has previously been used to identify *in vitro* budded MDVs^21^, and Ndufa10 and Mfn2, which have been shown to be regulated by PINK1 and Parkin, respectively^23–26^. We observed that MDVs floated to the 40/50% sucrose interface, as indicated by the presence of Uqcrfs1, Ndufa10 and Mfn2 in these fractions (Figure 1C & Figure S2B), which reflects the cargo selectivity of MDVs^27^ In contrast, non-mitochondrial proteins such as the lysosomal protein Lamp2a, the ER protein Calnexin and the synaptic protein Snap25 were either not present within the sucrose gradient or not specifically enriched at the 40/50% interface (Figure 1C & Figure S2B). Importantly, in the absence of cytosol (S), the intensity of mitochondrial proteins Uqcrfs1 and Vdac1 at the 40/50% sucrose interface was significantly lower, suggesting that MDV formation in this system is an active process that requires factors in the cytosol (Figure S2A). Ultrastructural examination of species present at the 40/50% sucrose interface using transmission electron microscopy revealed vesicular structures ranging from 40 nm to 240 nm in diameter, with a mean diameter of 120 nm, consistent with MDVs (Figure 1D). No difference in MDV size was observed between different PINK1 or Parkin genotypes, nor following antimycin A treatment (Figures 1D & S1B). These data indicate that the budding assay specifically generates MDVs that can be purified with minimal contamination from other cellular compartments.

We used western blots as a readout to investigate whether we could detect differences in MDV cargo following treatment with antimycin A or in PINK1^-/-^ or Parkin^-/-^ animals. Treatment with antimycin A, a mitochondrial complex III inhibitor, stimulates superoxide radical production and potentiates the formation of PINK1/Parkin-dependent MDVs^6, 21^. As a control, we incubated budding reactions with 0.1% DMSO (Figure 1A). While there was a trend towards greater inclusion of Uqcrfs1, Ndufa10 and Mfn2 in MDVs in response to antimycin A treatment, no significant difference was observed (Figure S2C&D). Furthermore, no difference was observed between wild-type and PINK1^-/-^ or Parkin^-/-^ animals (Figure S2C&D). Together, these data show that the *in vitro* budding assay provides a robust system to study brain MDVs and prompted us to adopt an unbiased approach to identify the cargo and machinery involved.

### Proteomic profiling of MDVs in brain reveal enrichment in OXPHOS proteins

To identify proteins enriched in MDVs, we isolated the 40/50% interfaces of sucrose gradients and subjected these fractions to bottom-up proteomics using ion mobility mass spectrometry. We compared the proteome of MDVs with their parental mitochondria in two independent sets of experiments using wild-type mouse brain (Data S1 (Data sets 1 and 2); N=3 biological replicates for each set of experiments, 2-3 animals per replicate).

A total of 4,367 proteins (Data set 1) and 4,796 proteins (Data set 2) were identified in the mitochondrial samples. Of those, 799 (Data set 1) and 862 (Data set 2) proteins were mitochondrial, representing 69% and 74% of the MitoCarta mitochondrial proteome, respectively^14^ A total of 991 proteins (Data set 1) and 2,328 proteins (Data set 2) were identified in the MDV samples. The large difference in the number of MDV proteins identified between the data sets may be due to the low abundance of the MDV samples, which leaves little margin of error for variation in handling of the samples.

To account for the variation in the low abundance MDVs and loading amounts between samples, we normalized LFQ values by expressing each LFQ value as a percentage of the total LFQ intensity for all proteins detected within that sample (% total LFQ, Data S1). We used Perseus to assess the differences between MDV and mitochondrial samples in proteins that had valid values in at least 70% of samples and at least 2 unique peptides. Statistical significance was determined by multiple t-testing visualized in a volcano plot, which identified MDV enriched and mitochondria enriched proteins (Figure 2A & S3A).

**Figure 2.**
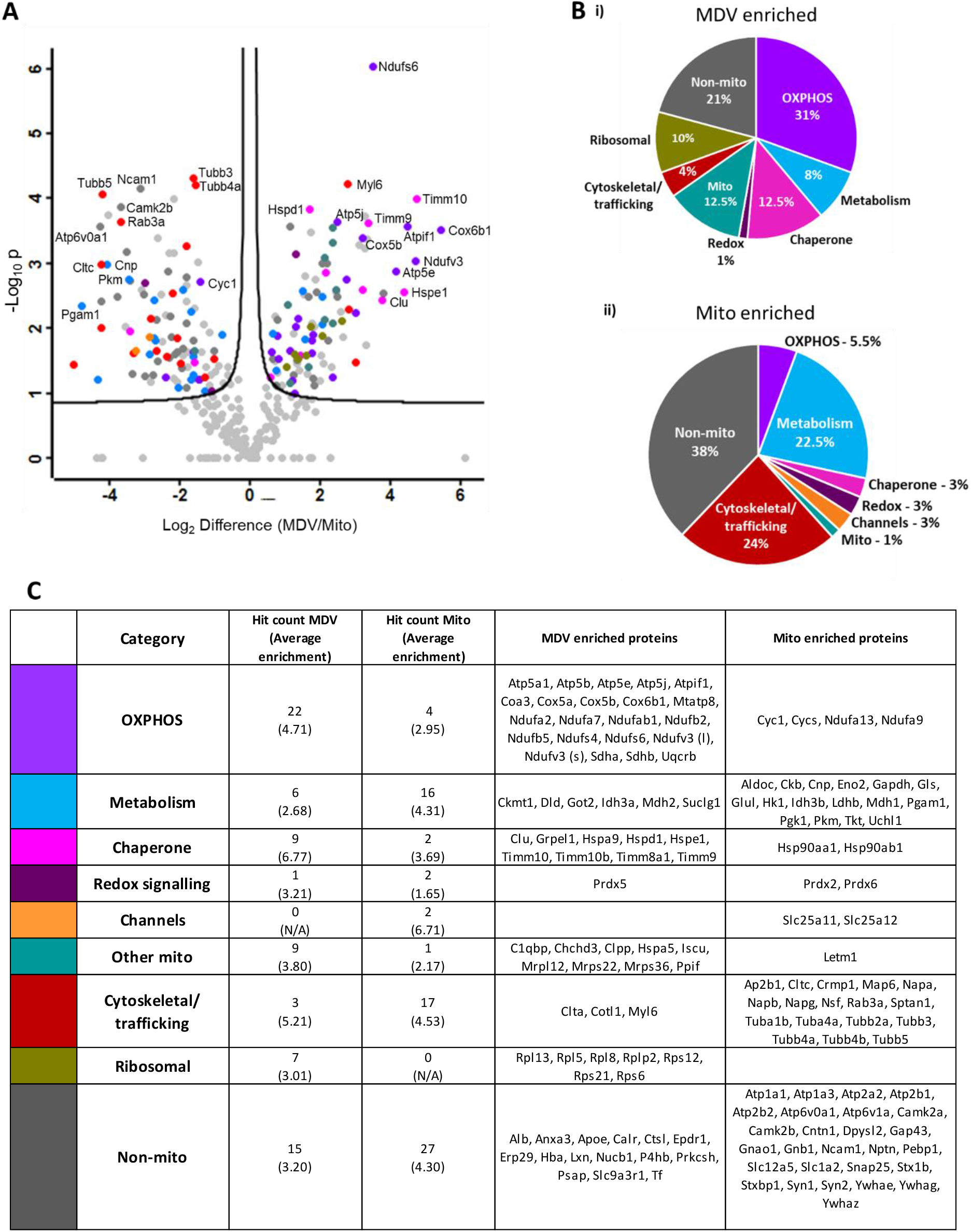
Proteomic identification of MDV enriched proteins. A) Volcano plot showing proteins significantly enriched (right hand side) or deenriched (left hand side) in MDVs compared with parental mitochondria in Data set 1. N=3. Coloured dots are hits that were also significant in Data set 2 (N=3, Figure S3A) and are coloured according to their functional assignment, as indicated in B&C. B) Pie charts showing the breakdown of hits according to their functional assignment in MDV enriched (i) or mitochondria enriched (ii) proteins. C) Table listing all hits and their functional assignment.

In the first set of experiments (N=3, Figure 2A), we identified 95 proteins as MDV enriched and 102 proteins as mitochondria enriched (Data set 1, Data S2). In the second set of experiments (N=3, Figure S3A), we identified 378 proteins as MDV enriched and 488 proteins as mitochondria enriched (Data set 2, Data S2). We took the overlap of hits from these two sets of experiments to be true hits and therefore counted 72 proteins as MDV enriched and 71 proteins as mitochondria enriched (Figure S3B). Of the proteins blotted for in Figure 1C, Uqcrfs1 and Ndufa10 were identified in MDVs, but were not significantly enriched in MDVs or mitochondria. Mfn2 was not detected in MDVs in Data Set 1, and though detected in MDVs in Data Set 2, it was defined as mitochondria enriched. Therefore, only a subset of MDV proteins are enriched in MDVs, suggesting mechanisms for cargo selection and packaging.

We also investigated oxidative post-translational modifications, as MDVs have previously been shown to transport oxidized cargo^27^. We initially identified an increase in oxidized methionine residues in MDVs compared with their parental mitochondria (Figure S4A, Data S3). However, a similar increase in methionine oxidation was observed when the parental mitochondria were subjected to the same sucrose gradient protocol as MDVs, suggesting this experimental procedure introduces high levels of methionine oxidation as an artefact (Figure S4B, Data S4). Therefore, specific methodologies for the study of oxidative modifications may be necessary to investigate molecular signatures of oxidative damage in MDVs.

We assigned the MDV or mitochondria enriched hits to a functional category (Figure 2B & Figure 2C) and observed that MDVs were enriched in OXPHOS proteins (31% of hits), while metabolic enzymes were enriched in mitochondria (22.5% of hits). Chaperone proteins and ribosomal proteins were also enriched in MDV hits compared to mitochondrial hits (Figure 2B&C). It is unclear why so many ribosomal proteins are highly enriched in MDVs, but it may be due to the high abundance of ribosomes within cellular extracts relative to the low abundance MDV samples. Mitochondria appear to be enriched in non-mitochondrial and cytoskeletal proteins, which is unsurprising because HM is a crude mitochondrial fraction, known to be contaminated with ER, MAM and synaptosomes. However, we can be sure that the enrichment of OXPHOS proteins in MDVs is not an artefact caused by the crudeness of the HM preparation: when non-mitochondrial and cytoskeletal components are removed from the analysis, the enrichment of OXPHOS proteins is maintained (40% MDV enriched vs 15% mitochondria enriched).

PCA analysis confirmed the distinct proteomic profiles of mitochondria and MDVs in both sets of experiments (Figure 3Ai&ii). Component 1 contributed the most variance (64.7%-75.6%), which was accounted for in both sets of experiments by mitochondrial complex I subunit Ndufs6, the chaperone protein Hspe1, the small TIM chaperones Timm10 and Timm9 and the ATP synthase inhibitory factor, Atpif1, again demonstrating the enrichment in OXPHOS and chaperone proteins (Figure 3Bi&ii).

**Figure 3.**
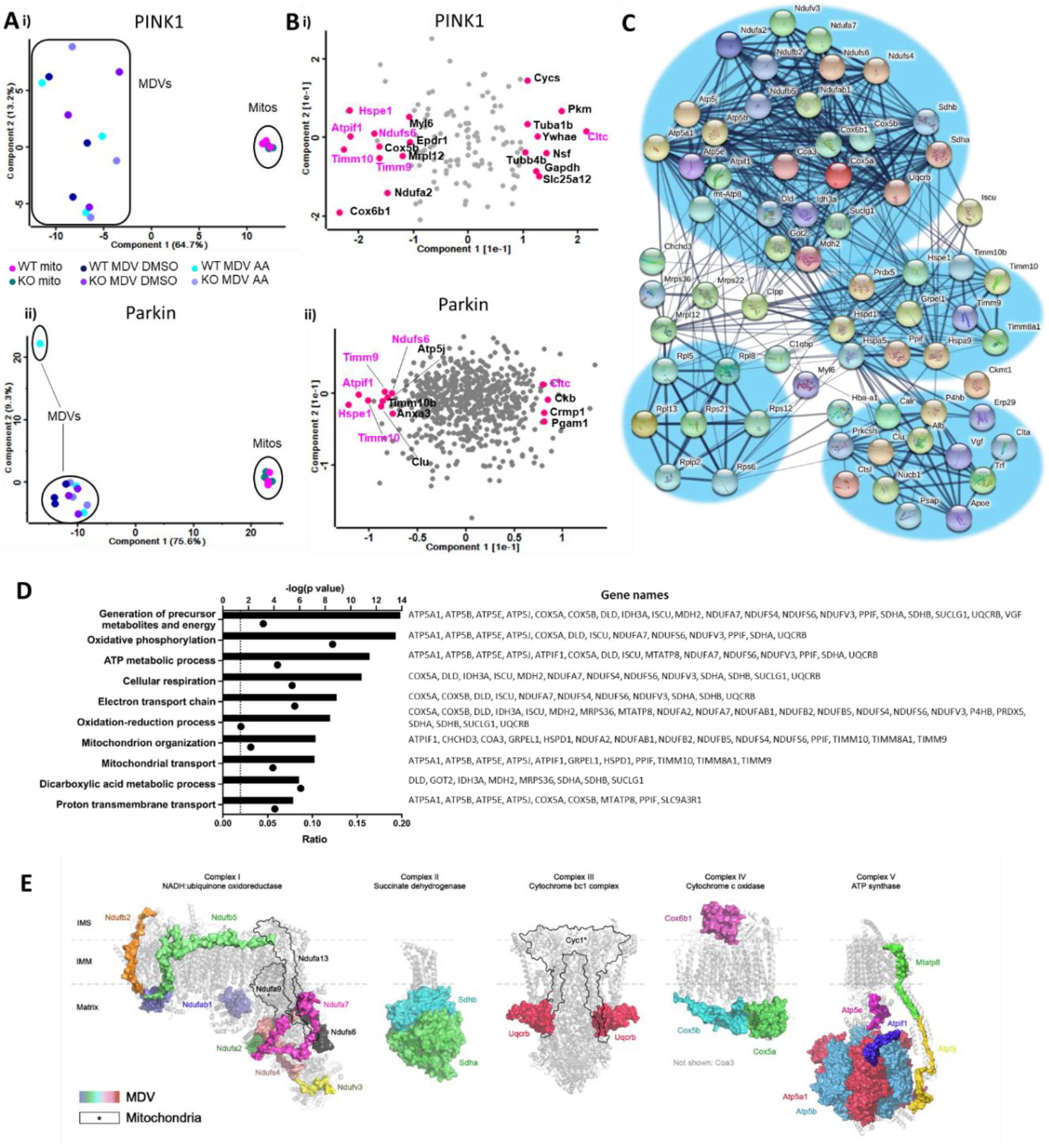
Cluster and pathway analysis of MDV cargo reveals enrichment in OXPHOS and small Tim chaperone proteins. A) PCA analysis demonstrates the distinct profiles of MDV and mitochondria along component 1 in both sets of experiments. B) PCA loading plot with proteins driving MDV (left) or mitochondrial (right) identity indicated in pink and labelled. C) STRING analysis on the 72 significantly enriched MDV proteins identified by the proteomic analysis revealed four clusters of proteins related to OXPHOS and metabolism (top cluster), translation (bottom left cluster), chaperone/redox proteins (middle cluster) and non-mitochondrial proteins (bottom right cluster). Only connected nodes are shown. The line thickness indicates the strength of the supporting data. D) Gene ontological analysis of MDV cargo reveals high enrichment of OXPHOS proteins. Bar graphs indicate the enrichment level of each pathway shown as −log(P-value). The corresponding dots indicate the ratio of identified proteins out of the total number of proteins in the pathway. E) OXPHOS proteins identified in the proteomic analysis are highlighted by their molecular surfaces in pyMOL 2.3.5. MDV enriched subunits are highlighted by their coloured molecular surface. Mitochondria enriched subunits are outlined in black and labelled with an asterisk (*). Each complex is depicted independent of respiratory supercomplexes in their lowest possible stoichiometry for simplicity. Complex I (human) PDB: 5XTD; complex II (bovine) PDB: 1ZOY; complex III (human) PDB: 5XTE; complex IV (human) PDB:5Z62; complex V (wild boar) PDB: 6J5J.

STRING analysis of the MDV enriched hits (Figure 3C) showed their clear partition into an OXPHOS and metabolic cluster (top), a chaperone cluster (middle), a ribosomal cluster (bottom left) and a non-mitochondrial cluster (bottom right). Gene ontological analysis further confirmed the enrichment of OXPHOS proteins MDVs (Figure 3D), with seven of the top ten gene ontological hits being related to oxidative phosphorylation. The gene ontological category ‘oxidative phosphorylation’ had the highest ratio of proteins in the pathway identified of any of the top ten categories (Figure 3D).

The clear enrichment of OXPHOS proteins in MDVs led us to question whether there were particular ‘hotspots’ in the complexes for MDV enrichment. Structural analysis revealed that several accessory subunits of the N module (Ndufa2, Ndufa7, Ndufs4, Ndufs6 & Ndufv3) and P_D_ module (Ndufab1, Ndufb2 & Ndufb5) of complex I were enriched in MDVs^28^ (Figure 3E). The matrix components of complex II (Sdha & Sdhb) and complex IV (Cox5a & Cox5b), as well as the complex III subunit Uqcrb were enriched in MDVs. Interestingly, Cox6b1, the subunit that bridges complex IV dimers together^29^, was enriched in MDVs. In complex V, 50% of the F1 subunits were detected as enriched in MDVs (Atp5a1, Atp5b & Atp5e), while the F0 subunits detected form a chain of interaction with the F1 head via Atp5j to Mtatp8. These observations may be indicative of new pathways for regulating assembly or disassembly of OXPHOS complexes.

### MDVs formed in the presence of antimycin A and/or the absence of PINK1 or Parkin are indistinguishable from those formed under basal conditions

We sought to identify cargo of MDVs stimulated by antimycin A and dependent upon the Parkinson’s disease linked proteins PINK1 and Parkin. Therefore, we compared the proteomic profiles of MDVs derived from mice null for PINK1 or Parkin with their wild-type littermates and in the presence of 50 μM antimycin A compared with the vehicle 0.1% DMSO. We observed no difference in any of the conditions, either by looking at the most significant hits from the wild-type analysis (Figure 4) or by directly comparing the profiles of wild-type and knockout or DMSO and antimycin A (Figure S5). These data suggest that PINK, Parkin and antimycin A dependent MDVs do not make up a large pool of MDVs in the brain and more specific approaches must be used to identify their unique cargo.

**Figure 4.**
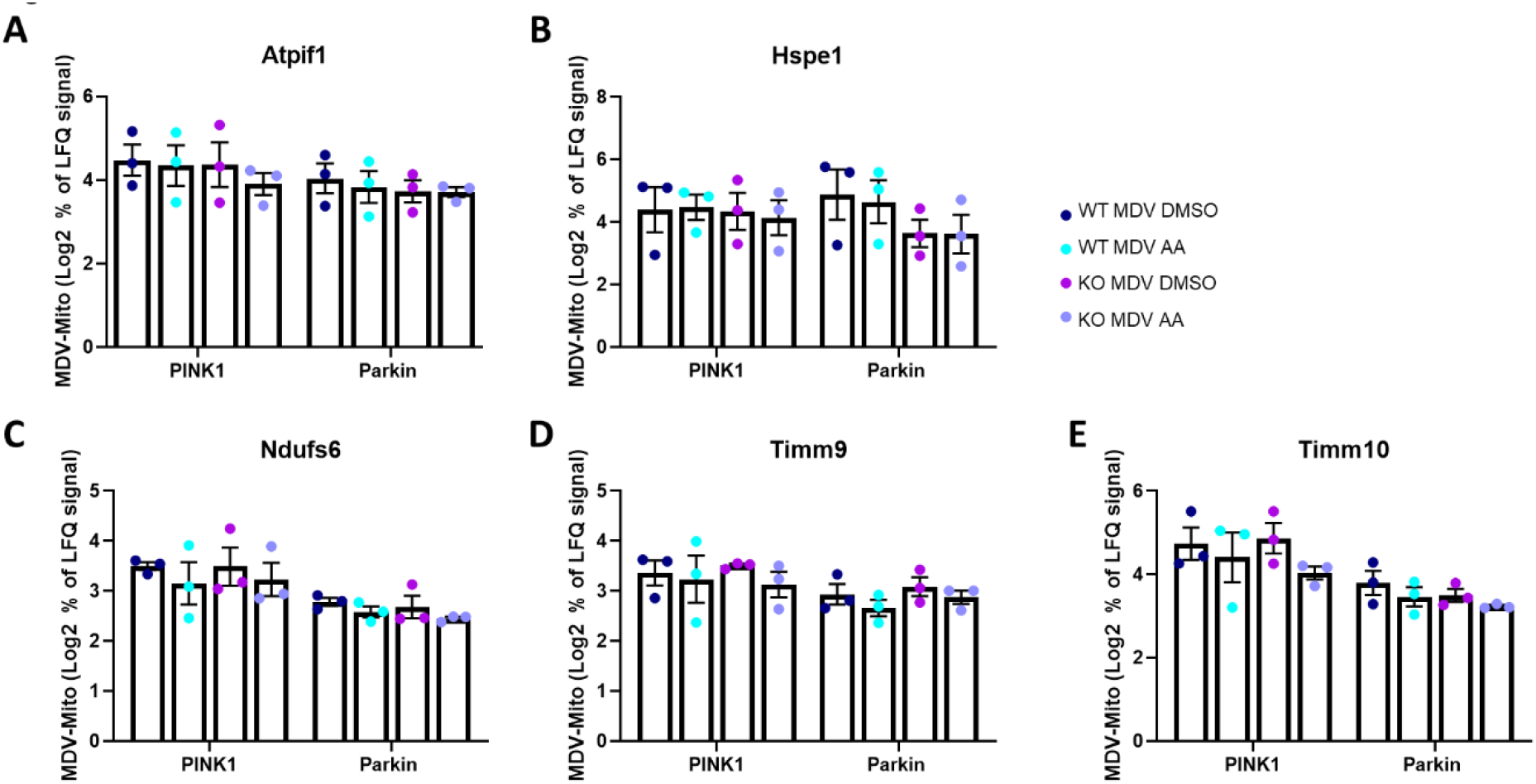
MDV enriched proteins are not dependent upon antimycin A, PINK1 or Parkin for MDV inclusion. The enrichment of top MDV hits from Figures 2&3, generated from wild-type mitochondria in the absence of oxidative stimulus (with DMSO), was compared with those generated in PINK1-null or Parkin-null animals and with stimulation by 50 μM antimycin A. No significant difference for any of the hits, including Atpif1 (A), Hspe1 (B), Ndufs6 (C), Timm9 (D) or Timm 10 (E), was observed. Two way ANOVA with Sidak’s multiple comparisons test.

## Discussion

In this study, we present for the first time an unbiased proteomic analysis of brain MDVs. Our results reveal the selective enrichment of a subset of OXPHOS proteins as MDV cargo. In particular, we demonstrate the inclusion of components of mitochondrial complex I and complex V in MDVs. Complex I and complex V components were previously thought to be unlikely to be incorporated into MDVs due to the size and complexity of the multi-protein complexes^6, 27^ In contrast, we found enrichment of a specific subset of components of all five electron transport chain complexes in MDVs. Importantly, we did not observe any difference in the enrichment of these subunits in response to antimycin A or in the absence of PINK1 or Parkin, suggesting a basal and general mechanism.

Interestingly, we observed that OXPHOS proteins identified as MDV enriched were localized to specific modules of each respiratory chain complex. The enrichment of subunits from a particular module may suggest the selective disassembly of particular areas of the complex in response to oxidative damage or for turnover of superfluous complexes. Alternatively, subassemblies that are deemed faulty during the process of complex assembly may be selected for inclusion in MDVs. Another attractive hypothesis is that subunits that are over-synthesized are removed for degradation via MDVs. SILAC experiments recently revealed the phenomenon of oversynthesis of unstable components and their subsequent degradation^30^. Complex I subunits were particularly prone to this phenomenon (27/33 subunits identified). This aligns with previous observations that complex I subunits have shorter half-lives compared to subunits from the other four electron transport chain complexes^31^. Of the nine complex I subunits we identified as enriched in MDVs, eight were identified as oversynthesized by Bogenhagen and Haley (all except Ndufab1). Given the additional role for Ndufab1 as an acyl carrier protein and in coordinating the assembly of respiratory complexes I, II, III and supercomplexes, it is possible that its inclusion in MDVs is related to this activity as opposed to its complex I activity^32^. Further investigations are warranted to determine whether turnover of over-synthesized complex I subunits is a function of MDVs.

Recent studies have also shown that some core subunits (NDUFS1, NDUFV1 and NDUFV2) of the matrix-facing N module of complex I are selectively turned over by the ClpXP protease in the mitochondrial matrix^33–34^ In light of this, it will be important to reconcile how quality control proteases crosstalk with MDV formation and cargo incorporation, particularly since Clpp was identified as enriched in MDVs in our study. It will also be necessary to assess whether and how MDVs regulate processed peptide efflux from mitochondria.

The enrichment of OXPHOS proteins in MDVs may suggest a quality control mechanism for the complexes central to the generation of ATP. This would align with the observation of Vincow et al, who hypothesized that their results showing the turnover of respiratory chain proteins in a Parkin, but not Atg7, dependent manner may indicate their inclusion in MDVs^35^. A more recent study from the same group calculated that only 1/3 of mitochondrial protein turnover occurs via autophagy, suggesting that pathways like MDVs or proteasomal degradation play a significant role in this regard^36^. Here we provide evidence that MDVs are indeed enriched in respiratory chain proteins in a mammalian system. Four of the OXPHOS proteins we identified as enriched in MDVs were also identified in Vincow et al’s study (Ndufa7, Sdhb, Uqcrb, Cox5a). However, we did not observe that their inclusion in MDVs was dependent on Parkin. This may be a species difference, as their study was in Drosophila while ours was in mouse. As previously mentioned, it is also possible that the PINK/Parkin vesicles are more specialized and less abundant and therefore not significantly detected in this bulk analysis of MDVs. It is also important to note that in this assay, we were unable to determine the cellular fate of the MDVs detected. Further cellular and functional studies will be required to elucidate the mechanism for and consequences of OXPHOS selectivity in MDVs.

Another striking observation was the enrichment of the small TIM chaperones in MDVs (Timm10, Timm10b, Timm9, Timm8a1). The small TIM proteins localize to the intermembrane space where, as heterohexameric complexes of Timm9/Timm10 or Timm8/Timm13, they chaperone membrane proteins between the TOM complex and the TIM22 complex, for multipass inner mitochondrial membrane proteins, or the SAM complex, for outer mitochondrial membrane beta-barrel proteins^37^ Very few of their client proteins are known, but in yeast include Tim22, Tim23, Tim17, Mic17 and Cox12 (Cox6b1)^38^. More recent investigations have identified its mammalian client proteins, such as the metabolite carriers Slc25a3, Slc25a4, Mpc1 and Mpc2^39–41^. Cox6b1 was the only putative small Tim protein cargo identified as enriched in MDVs, suggesting that the inclusion of the small Tim proteins in MDVs may not be related to faulty chaperoning of their cargo.

Timm9, Timm10 and Timm10b were previously identified in a screen for mitophagy regulators^42^. While Hoshino and colleagues did not investigate the mechanism through which the small TIM proteins promote mitophagy, our identification of the small TIMs, along with other known regulators of mitophagy such as Atpif1^43^, as MDV enriched, raises interesting possibilities. One exciting hypothesis is that when mild damage occurs, these proteins are selectively removed from mitochondria so as not to induce wholesale mitophagy. In this scenario, MDVs both remove mild damage and ensure that over-reaction to damage does not occur.

In conclusion, we have described the proteomic profile of MDVs for the first time, deepening our understanding of this biological cellular pathway. Functional studies will be required to tease apart which cargo are unique to particular subsets of MDVs and how they are regulated and trafficked. We believe that this dataset is an exciting starting point from which to investigate these questions and reveals further complexity and novel cargo in MDVs.

## Acknowledgements

This work was supported by a Foundation grant from CIHR (FDN-154301) and a Canada Research Chair (Tier 1) in Parkinson’s disease to EAF. RFR was supported by a Parkinson’s Foundation Post-Doctoral Fellowship (PDF-FBS-1638). ANB is supported by a CIHR doctoral award. JFT is supported by a Canada Research Chair (Tier 2) in Structural Pharmacology. We thank Dr. Esther del Cid Pellitero and Frédérique Larroquette for mouse colony care and management. We also thank Jeannie Mui at the Facility for Electron Microscopy Research of McGill University for help with TEM sample processing, microscope operation and for technical advice. We are grateful to Dr. Benoit Vanderperre (McGill University) and Dr. Brent Ryan (University of Oxford) for productive discussions about data analysis and interpretation for this manuscript.

## Supporting Information

Figure S1: MDVs derived from PINK1 or Parkin null animals are indistinguishable from those formed from wild type animals.

Figure S2: Quantification of protein enrichment in MDVs by western blot.

Figure S3: Proteomic identification of MDV enriched proteins.

Figure S4: Analysis of oxidative post-translational modifications in MDVs and their parental mitochondria.

Figure S5: MDV enriched proteins are not dependent upon antimycin A, PINK1 or Parkin for MDV inclusion.

Data S1: MaxQuant Protein Groups search outputs for two sets of independent experiments (N=3 each).

Data S2: Proteins identified as significantly enriched in Perseus statistical analysis.

Data S3: Analysis of post-translational modifications in mitochondria and MDV samples.

Data S4: Analysis of post-translational modifications in mitochondria following subjection to sucrose gradient preparation.

