## Supplementary Figures for "Proteomic profiling of mitochondrial-derived vesicles in brain reveals enrichment of respiratory complex sub-assemblies and small TIM chaperones"

Table of contents

Figure S1: MDVs derived from PINK1 or Parkin null animals are indistinguishable from those formed from wild type animals.

Figure S2: Quantification of protein enrichment in MDVs by western blot.

Figure S3: Proteomic identification of MDV enriched proteins.

Figure S4: Analysis of oxidative post-translational modifications in MDVs and their parental mitochondria.

Figure S5: MDV enriched proteins are not dependent upon antimycin A, PINK1 or Parkin for MDV inclusion.

Data S1: MaxQuant Protein Groups search outputs for two sets of independent experiments (N=3 each).

Data S2: Proteins identified as significantly enriched in Perseus statistical analysis.

Data S3: Analysis of post-translational modifications in mitochondria and MDV samples.

Data S4: Analysis of post-translational modifications in mitochondria following subjection to sucrose gradient preparation.

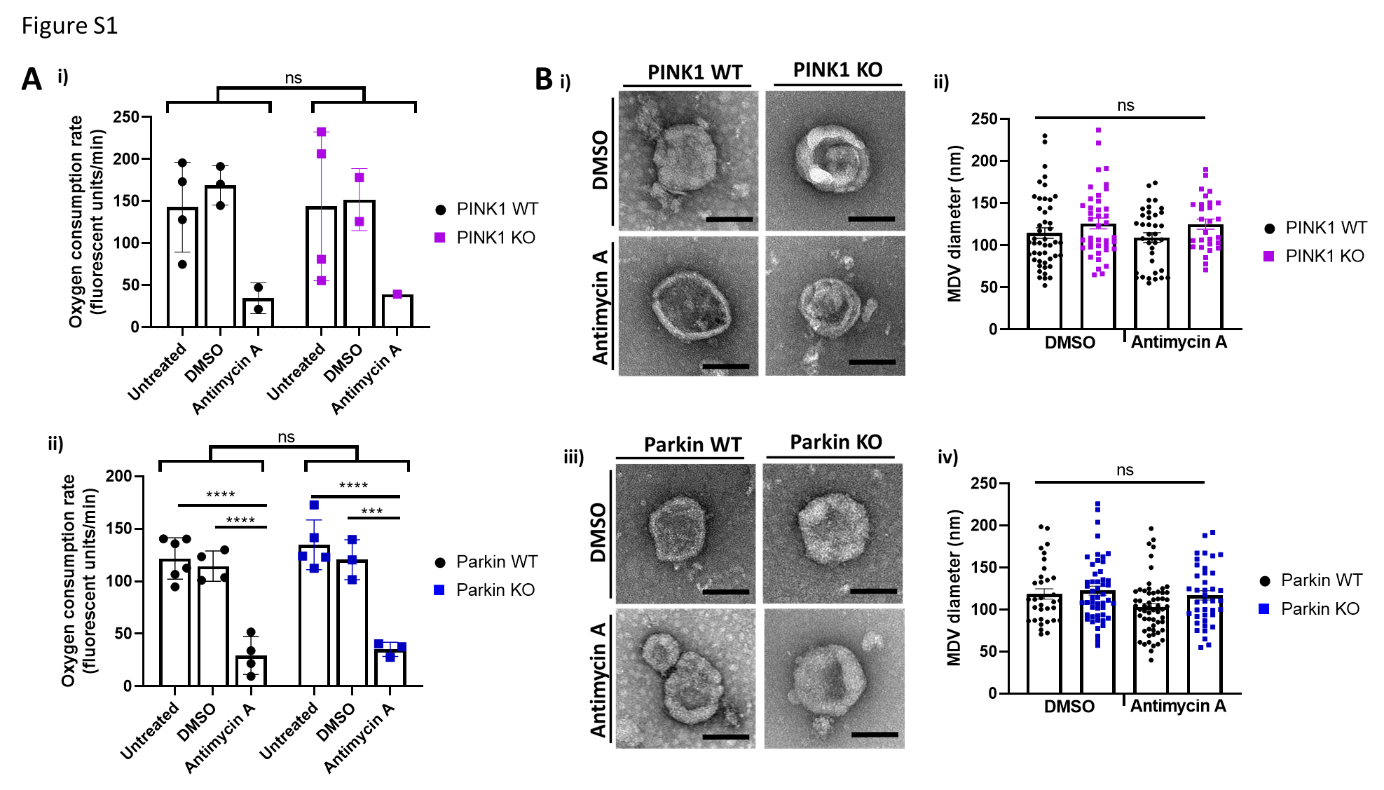
**Figure S1.** MDVs derived from PINK1 or Parkin null animals are indistinguishable from those formed from wild type animals. A) Mitochondria (HM) isolated from wild-type, PINK1 null or Parkin null mouse brain are metabolically active and responded to the complex III inhibitor antimycin A (AA), as assessed by MitoXpress respiratory assay. No difference between wilde-type and knock-out mice was detected. Two way ANOVA with Tukey’s multiple comparisons test. ****: p<0.0001. B) i&iii) Transmission electron micrographs of MDVs stained with uranyl acetate. Scale bar: 100nm. ii&iv) Quantification of MDV diameter from EM images. No significant differences between wild-type or null mice, or MDVs formed in the presence of DMSO or antimycin A were observed. Two way ANOVA with Sidak’s multiple comparison test.

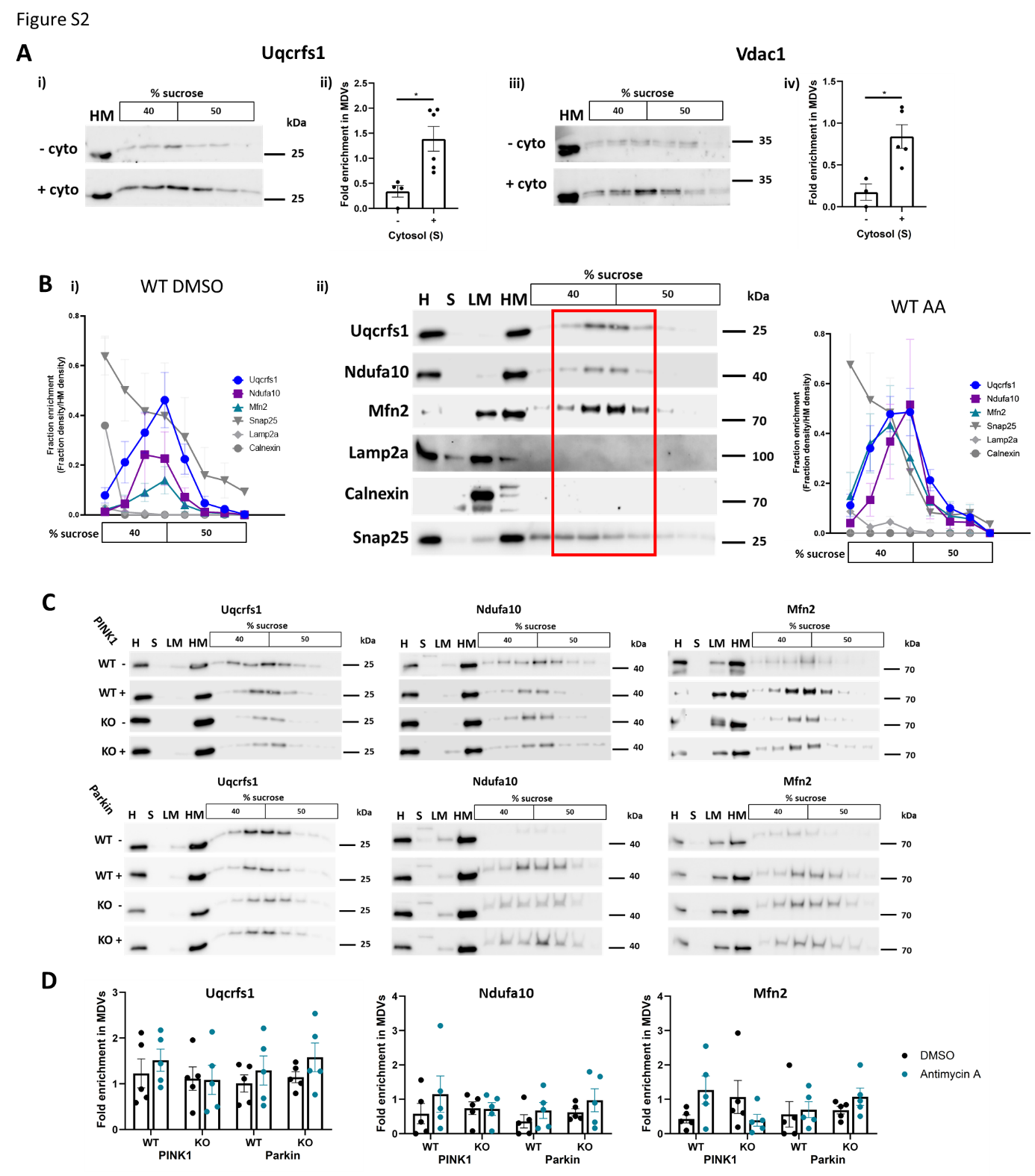

**Figure S2.** Quantification of protein enrichment in MDVs by western blot. A) MDV generation is stimulated by including cytosol in the budding reaction. Representative blots for Uqcrfs1 (i) and Vdac1 (iii) and their quantifications (ii&iv) are shown. B) The enrichment in each sucrose gradient fraction of mitochondrial (Uqcrfs1, Ndufa10, Mfn2) and non-mitochondrial (Snap25, Lamp2a, Calnexin) were assessed in the presence of DMSO (i, quantification with error bars of representative blots in Figure 1C) or antimycin A (ii&iii) as in Figure 1C. C) The enrichment of Uqcrfs1, Ndufa10 and Mfn2 in MDVs (fractions 2-5, compared with HM density) were assessed in wild-type mice and their knockout littermates (PINK1-/-, top panel; Parkin-/-, bottom panel) in the presence of DMSO or antimycin A. No significant difference between the conditions was observed. Two way ANOVA with Tukey’s multiple comparisons test.

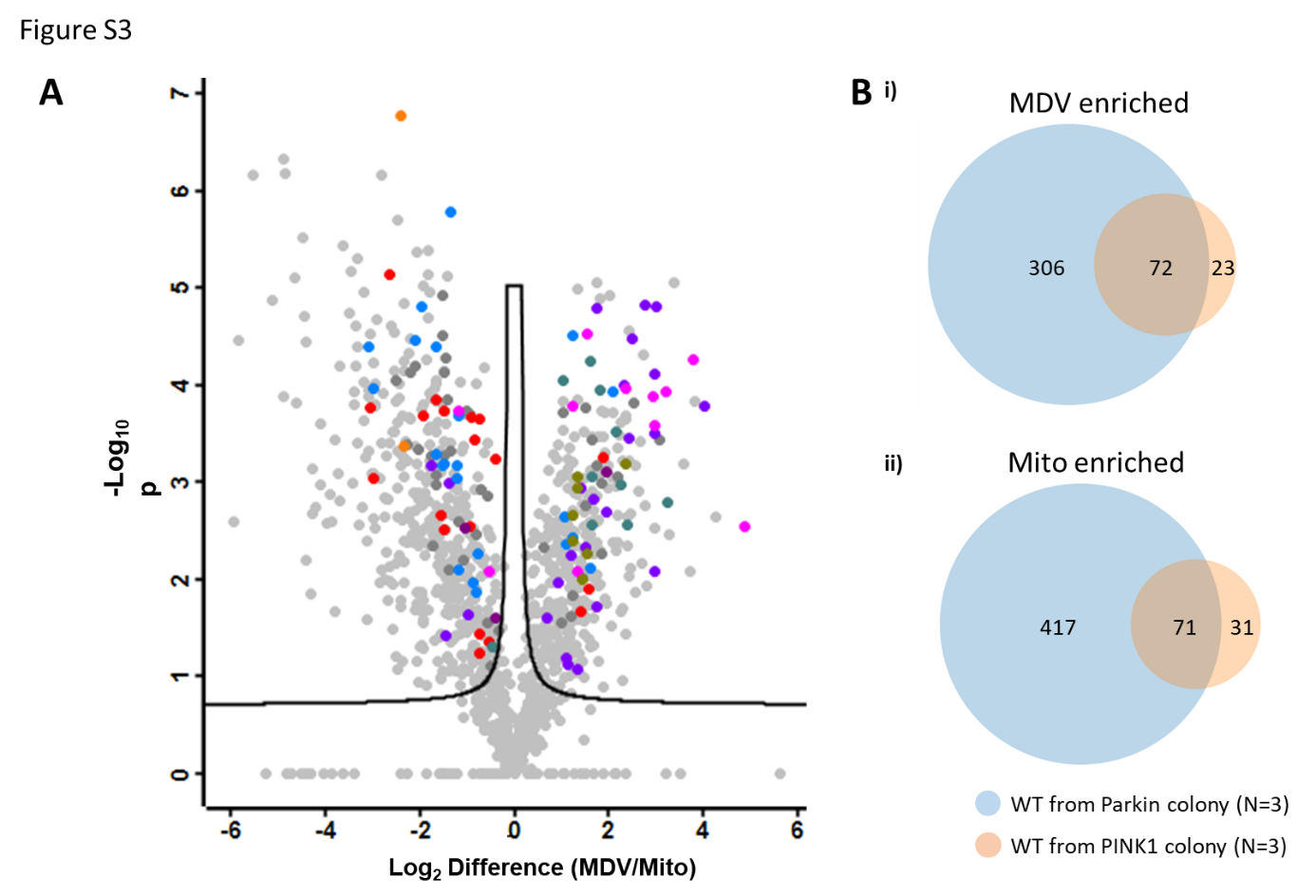

**Figure S3.** Proteomic identification of MDV enriched proteins. A) Volcano plot for the second set of experiments (Data set 2) showing proteins significantly enriched (right hand side) or deenriched (left hand side) in MDVs compared with parental mitochondria. N=3. Coloured dots are hits that were also significant in a separate experiment (N=3, Figure 2A) and are coloured according to their functional assignment, as indicated in Figure 2B&C. B) Number of hits in the two independent sets of experiments (N=3 for each) defined as MDV enriched (i) or mitochondria enriched (ii).

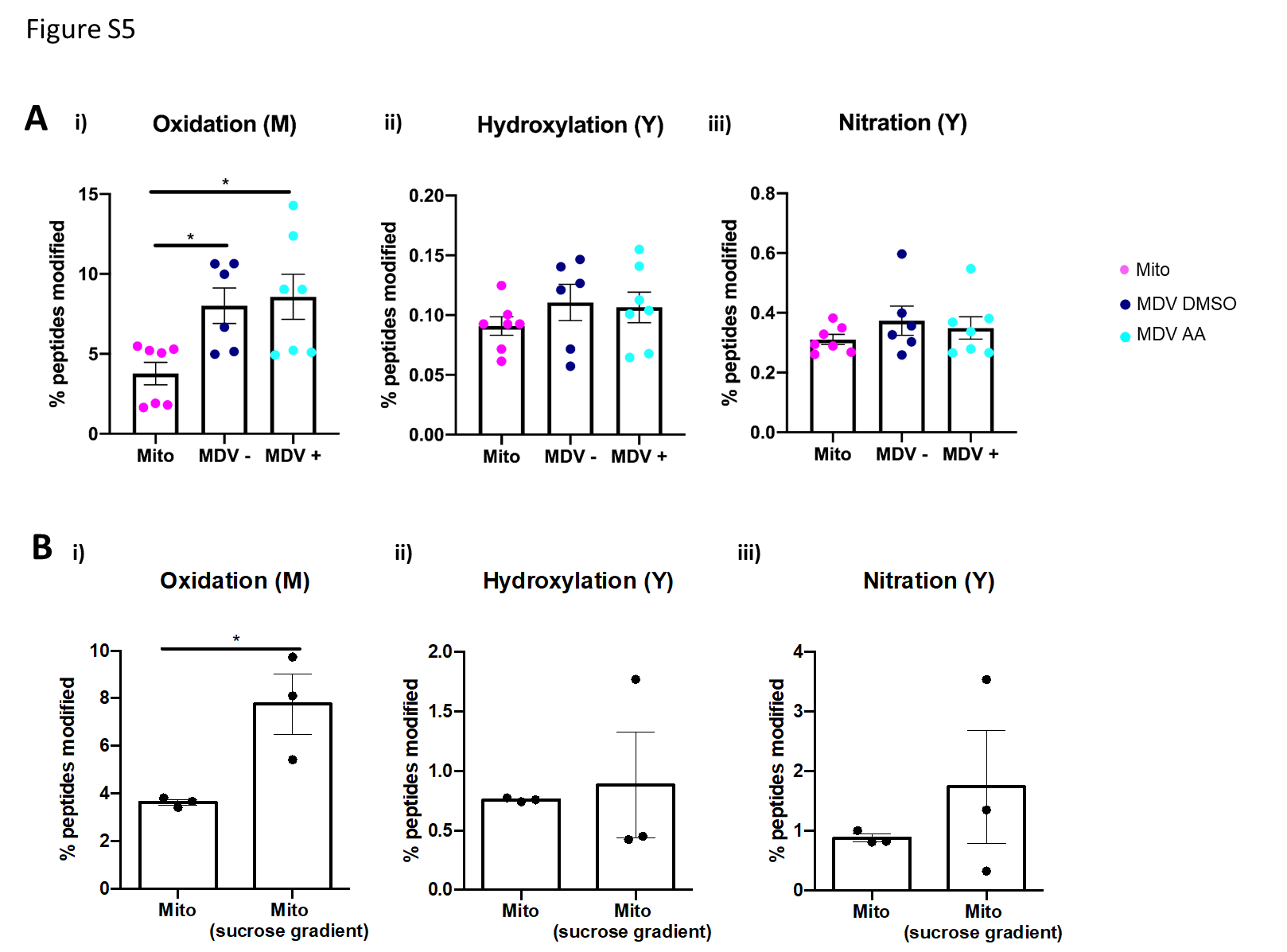

**Figure S4.** Analysis of oxidative post-translational modifications in MDVs and their parental mitochondria. A) Global levels of (i) methionine oxidation, (ii) tyrosine hydroxylation and (iii) tyrosine nitration were calculated in mitochondrial (mito) and MDV samples without (MDV-) or with (MDV+) antimycin A stimulation. One way ANOVA with Dunnett’s multiple comparisons test. *: p<0.05, **: p<0.01, ***: p<0.001. B) Global levels of (i) methionine oxidation, (ii) tyrosine hydroxylation and (iii) tyrosine nitration were calculated in mitochondrial samples before and after sucrose gradient fractionation. Unpaired two-tailed student’s t test. *: p<0.05.

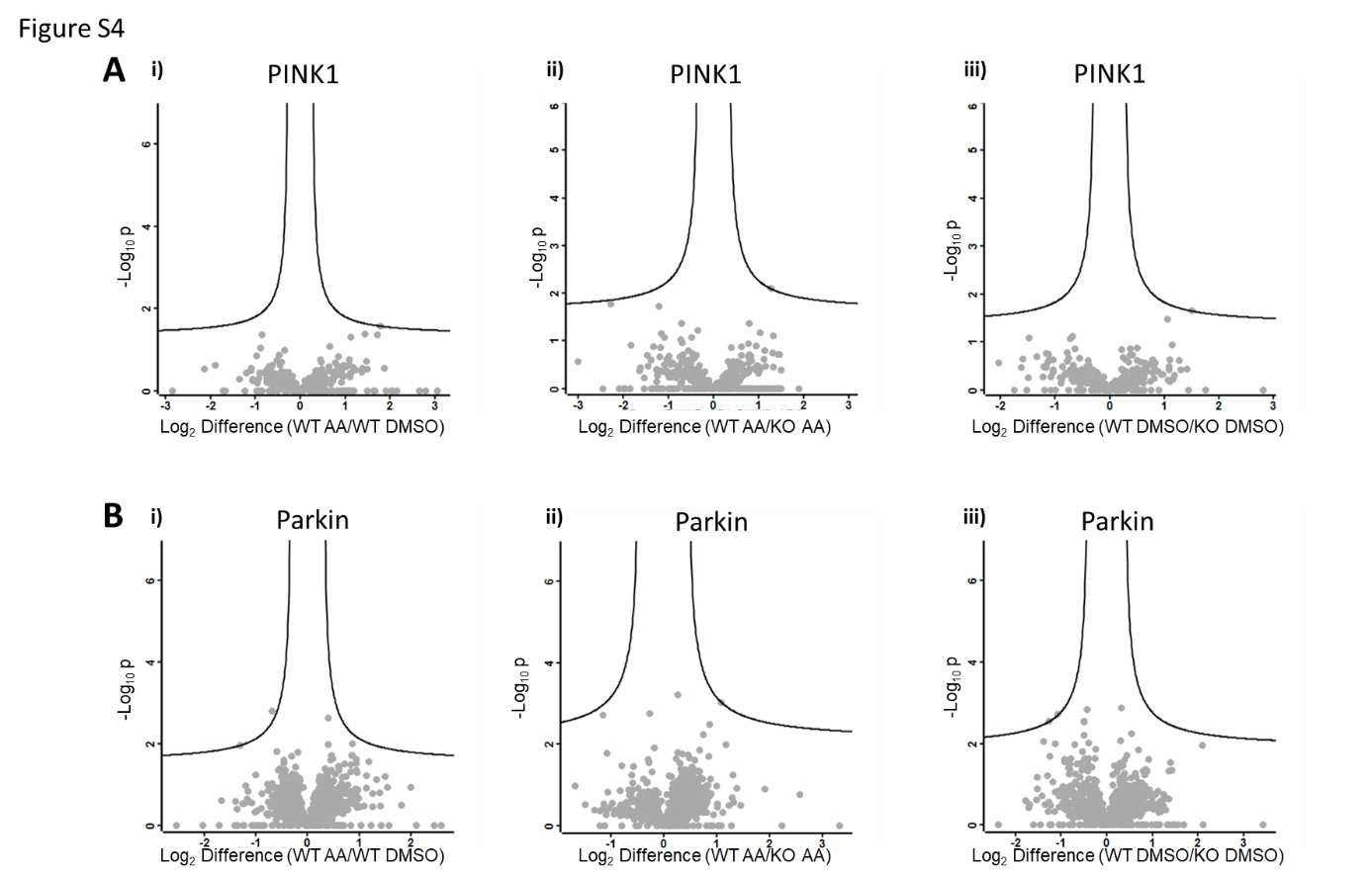

**Figure S5.** MDV enriched proteins are not dependent upon antimycin A, PINK1 or Parkin for MDV inclusion. Volcano plots from two independent sets of experiments (A&B, N=3 each) showing that no significant difference in MDV cargo is observed between MDVs formed in the presence of DMSO or antimycin A (Ai, Bi), or between wild type and PINK1-null animals (Aii, Aiii) or wild type and Parkin-null animals (Bii, Biii).
